# A defining member of the new cysteine-cradle family is an aECM protein signalling skin damage in *C. elegans*

**DOI:** 10.1101/2024.04.11.589058

**Authors:** Thomas Sonntag, Shizue Omi, Antonina Andreeva, Jeanne Eichelbrenner, Andrew D. Chisholm, Jordan D. Ward, Nathalie Pujol

## Abstract

Apical extracellular matrices (aECMs) act as crucial barriers, and communicate with the epidermis to trigger protective responses following injury or infection. In *Caenorhabditis elegans*, the skin aECM, the cuticle, is produced by the epidermis and is decorated with periodic circumferential furrows. We previously showed that mutants lacking cuticle furrows exhibit persistent immune activation (PIA). In a genetic suppressor screen, we identified *spia-1* as a key gene downstream of furrow collagens and upstream of immune signalling. *spia-1* expression oscillates during larval development, peaking between each moult together with patterning cuticular components. It encodes a secreted protein that localises to furrows. SPIA- 1 shares a novel cysteine-cradle domain with other aECM proteins. SPIA*-*1 mediates immune activation in response to furrow loss and is proposed to act as a sensor of cuticle damage. This research provides a molecular insight into intricate interplay between cuticle integrity and epidermal immune activation in *C. elegans*.

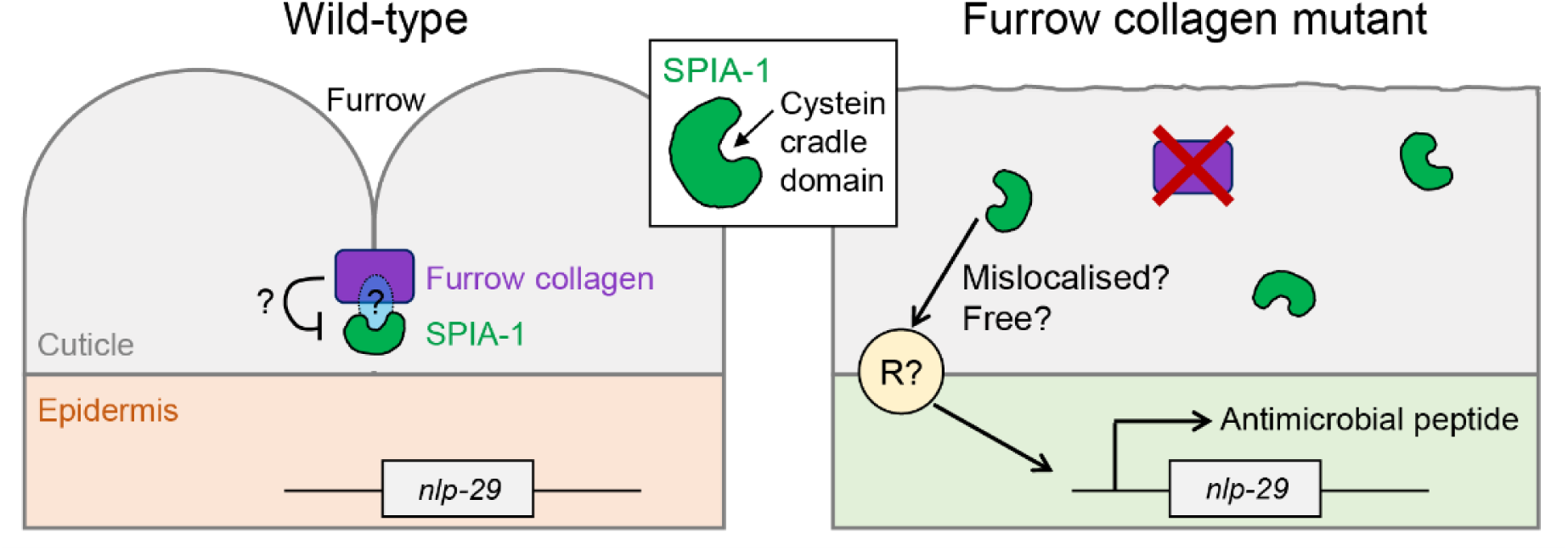

## Introduction

All multicellular organisms must protect themselves from injury and pathogens. *Caenorhabditis elegans* lacks an adaptive immune system and motile immune cells. Instead, it relies on its epithelial barriers to defend itself against environmental threats. This makes it a powerful model to address the question of how epithelial cells detect damage. In *C. elegans*, the skin is characterised by a rigid but flexible apical extracellular matrix (aECM), known as the cuticle, that surrounds a single syncytial epidermal layer (reviewed in (Sundaram and Pujol, 2024)). The cuticle surface contains circumferential-oriented furrows distributed periodically over the entire body length (Adams et al., 2023; Cox et al., 1981; McMahon et al., 2003). Embryos assemble the first larval cuticle, and then during each larval stage, a new cuticle is assembled, and the old one shed, in a process known as moulting (Lažetić and Fay, 2017). A transient precuticle is assembled to help to pattern the new cuticle and to shed the old one (Sundaram and Pujol, 2024).

The cuticle serves not only as a protective barrier against environmental insults but also as a dynamic interface that communicates crucial signals to the underlying epidermal tissue. We have previously described how cuticle damage triggers a series of responses in the epidermis. These responses can be set off by physical injury, infection with the fungus *Drechmeria coniospora*, or during the cyclic process of moulting. The organism’s ability to mount a protective transcriptional response maintains tissue integrity to combat potential threats (Martineau et al., 2021; Pujol et al., 2008a; Sundaram and Pujol, 2024).

Mutants lacking periodic furrows have emerged as a valuable model for studying the interplay between cuticle integrity and epidermal immune activation. Mutations in any of the six furrow collagens (DPY-2, DPY-3, DPY-7, DPY-8, DPY-9, DPY-10) lead to the absence of periodic furrows in the cuticle (Cox et al., 1980; McMahon et al., 2003; Thein et al., 2003). We have previously shown that the same mutations exhibit a persistent immune activation (PIA), similar to the response triggered by moulting, physical injury or skin infection. This immune response involves the activation of the pivotal p38 MAPK/PMK-1 signalling pathway and the downstream SNF-12/SLC6 transporter and STAT-like transcription factor STA-2 (Dierking et al., 2011; Dodd et al., 2018; Pujol et al., 2008b). During infection or injury, the most upstream components known are the Damage Associated Molecular Pattern (DAMP) receptor DCAR-1, a GPCR, and the Gα protein GPA-12 (Zugasti et al., 2014). While loss of STA-2 or SNF-12 fully abrogates the induction of an immune response in furrow collagen mutants, inactivation of DCAR-1 only reduces it partially (Zugasti et al., 2014). We therefore proposed that a parallel mechanism must link the monitoring of furrow collagens’ integrity to the activation of the immune response in the epidermis.

To gain deeper insights into how cuticle damage is sensed by the epidermis, we conducted a targeted genetic suppressor screen to identify genes acting downstream of furrow collagens and upstream of, or in parallel to, GPA-12. Notably, one suppressor identified in this screen harbours a mutation in the gene *spia-1* (*Suppressor of Persistent Immune Activation*). This gene encodes a small nematode-specific secreted protein sharing a C-terminal domain with five other *C. elegans* proteins, including DPY-6, a mucin-type protein with a conserved role in cuticle deposition (Sun et al., 2022). The structured core of this common C-terminal domain (named CCD-aECM) is formed by conserved cysteine interactions predicted to allow potential homomeric and heteromeric interactions. Our expression and genetic analyses suggest that SPIA-1 functions as a secreted aECM protein, localised to the furrows, potentially relaying information about the state of the furrow collagens to the underlying epidermis.

## Results & Discussion

### Identification of *spia-1* as a suppressor of a constitutive immune response

We previously showed that wounding and infection of *C. elegans* trigger an immune response, characterised by the induction of the expression of the antimicrobial peptide (AMP) gene *nlp-29* in the epidermis (Couillault et al., 2004; Martineau et al., 2021; Pujol et al., 2008a; Taffoni et al., 2020; Zugasti et al., 2014). Interestingly, mutants in furrow collagens, which lack the organised circumferential furrow structure (“furrow-less mutants”) (Aggad et al., 2023), also have a persistent immune activation (PIA) (Pujol et al., 2008b; Zugasti et al., 2014; Zugasti et al., 2016), in parallel to constitutively active detoxification and hyperosmotic responses (Dodd et al., 2018). The fact that these 3 responses are induced by the absence of furrows, yet differ in their signalling and effectors, led to the suggestion that a cuticle-associated damage sensor coordinates these 3 responses (Dodd et al., 2018).

To characterise this potential damage sensing mechanism, we conducted a genetic suppressor screen, designed to identify the upstream components of the pathway leading to the induction of the immune response in furrow-less mutants. The screen relies on the observation that a constitutively active form of GPA-12 (GPA-12gf) provokes a PIA by activating the p38/PMK-1 – STA-2 pathway, and on the use of a conditional promoter that is only active in the adult epidermis (Lee et al., 2018). In addition to a construct expressing GPA*-*12gf uniquely in the adult epidermis, the strain IG1389 we designed has the well- characterised *frIs7* transgene containing the *nlp-29* promoter driving GFP expression (*nlp-29*p::GFP) and a control DsRed transgene constitutively expressed in the epidermis, providing an internal control for the integrity of the epidermis and nonspecific transgene silencing (Pujol et al., 2008a). In the IG1389 strain, the *nlp-29*p::GFP reporter is not expressed in larvae but constitutively expressed in the adult, due to the expression of GPA-12gf ((Ziegler et al., 2009); Fig 1A). When any of the six furrow collagen genes, including *dpy-7*, is inactivated by RNAi in this strain, worms exhibit a high level of GFP at all developmental stages (green larvae-green adults) (Fig 1A, 1B and S1A). Inactivating any gene acting downstream of GPA*-*12, like *sta-2*, completely abolishes the expression of *nlp-29*p::GFP at all stages (red larvae-red adults; the so-called “no induction of peptide after infection” (Nipi) phenotype (Pujol et al., 2008a)) (Fig 1B and S1A). However, if the inactivated gene acts upstream of, or in parallel to, GPA-12, the expression of *nlp-29*p::GFP should be suppressed in larvae but reactivated by GPA-12gf in the adult (Fig S1A), as observed with *dcar-1,* which acts upstream of *gpa-12* (Fig 1B).

**Fig 1.**
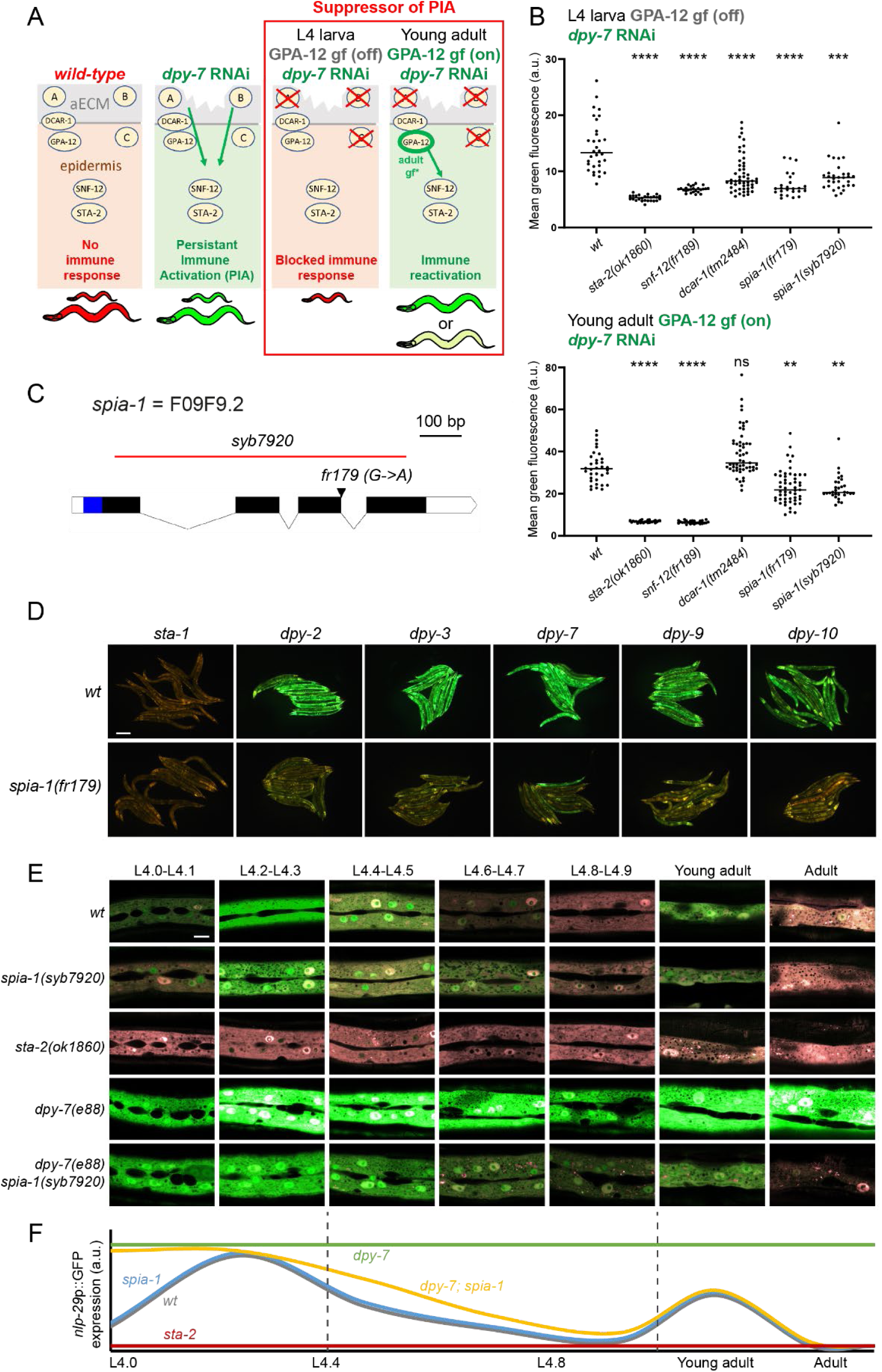
Loss of *spia-1* suppresses furrow collagen AMP induction. (A) Design of the suppressor screen. The strain carries the *frIs7* transgene, containing an AMP transcriptional reporter (*nlp-29*p::GFP) and a control transgene (*col-12*p::DsRed) constitutively expressed in the epidermis. Under standard growth conditions, worms only express the control transgene and are red at all stages (left). RNAi inactivation of any furrow collagen gene, like *dpy-7*, leads to the expression of *nlp-29*p::GFP in a PMK-1/STA-2 dependent manner: worms appear “green” at all stages (middle). The strain used for the suppressor screen additionally bears the *frIs30* construct to express a gain of function of GPA-12 in the epidermis, only from the young adult stage (*col-19*p::GPA-12gf). In this strain, inactivation of a gene downstream of GPA-12 eliminates the expression of *nlp-29*p::GFP in both larvae and adults (Nipi phenotype (Pujol et al., 2008a)), whereas inactivation of any gene acting upstream of, or in parallel to, GPA-12, inhibits the expression of *nlp-29*p::GFP in the larvae but not in the adult due to the activation of GPA*-*12 (red larvae, green adults, right). This rescue is total if the targeted gene acts upstream of GPA-12 (A, *dcar-1*), but only partial if it acts in parallel to GPA-12 (B/C). (B) Quantification of the green fluorescence in worms carrying *frIs7* and *frIs30* constructs in different mutant backgrounds, upon *dpy-7* RNAi, in L4 and young adult stages (n>25). Statistics were made by comparing to the corresponding *wt* control; \*\**p* < 0.01; \*\*\**p* < 0.001; \*\*\*\**p* < 0.0001. (C) Structure of the *spia-1* genomic locus. The location of the *fr179* mutation is indicated with an arrowhead, the extent of the *syb7920* deletion is shown with a red line. Exons are shown as black boxes, introns as solid lines. UTR are represented as white boxes; the blue region shows the sequence encoding the signal peptide of *spia-1*. (D) *spia-1(fr179)* suppresses the induction of *nlp-29*p::GFP in young adult worms after RNAi inactivation of furrow collagen genes. Wild-type and *spia-1(fr179)* mutants carrying the *frIs7* transgene were treated with the indicated RNAi bacteria, with *sta-1* used as a control (see Mat&Methods). Red and green fluorescences were visualised simultaneously in all images. Representative images of young adults from one of three experiments are shown; scale bar, 200 µm; see Fig S1C and S1D for quantification with the COPAS Biosort. (E) Oscillation of *nlp-29*p::GFP expression from L4 to adulthood. Representative confocal images of different mutant strains carrying the *frIs7* transgene, red and green fluorescences were visualised simultaneously. The L4 stage is subdivided into sub-stages with the shape of the vulva (Mok et al., 2015); n>5, scale bar, 10 µm. (F) Proposed schematic illustration of *nlp-29*p::GFP oscillation shown in E, units are arbitrary and not to scale.

We mutagenised the strain IG1389 using ethyl methanesulfonate (EMS), then transferred synchronised F2 progeny onto *dpy-7* RNAi plates at the L1 stage and screened for mutants that suppressed the PIA phenotype. Many mutants corresponded to genes acting downstream of *gpa-12*, as they blocked the PIA at both larval and adult stages. Complementation tests allowed us to identify new alleles of components of the known pathway, including *snf-12(fr189)*. Interestingly, another subset of mutants had a phenotype comparable to *dcar-1*, i.e the expression of *nlp-29*p::GFP was suppressed in larvae but reactivated in the adult. Among them, the *fr179* mutant had the clearest phenotype (Fig 1B). We called the *fr179* mutant *spia-1* (*Suppressor of Persistent Immune Activation).* We confirmed that in the absence of *dpy-7* RNAi, *spia-1(fr179)* mutation also did not suppress the *gpa-12*-induced PIA observed in adults, while *sta-2(ok1860)* completely abrogated it (Fig S1B). This data suggested that *spia-1* does not act downstream of *gpa-12*. Moreover, unlike *dcar-1* mutation, *spia-1(fr179)* still partly blocked the PIA in adults, when it is provoked by a combination of *dpy-7* inactivation and *GPA-12gf* gain of function (Fig 1B). Together, these data suggest that SPIA-1 acts in a partially non-redundant pathway parallel to GPA-12.

We backcrossed the *spia-1(fr179)* strain relying on the suppression of *nlp-29*p::GFP induction upon *dpy-7* RNAi for the selection of *spia-1(fr179)* progeny. The underlying molecular lesion was characterised by mapping through whole genome sequencing (WGS) of a pool of backcrossed *spia-1(fr179)* independent recombinant mutants (Doitsidou et al., 2010; Labed et al., 2012). The *spia-1(fr179)* worms carry a mutation in a splice donor site of the gene *F09F9.2* (hereafter *spia-1*), predicted to result in a transcript with a frameshift and the introduction of a premature stop codon leading to a truncated protein of 133 aa (Fig 1C). We generated by CRISPR the allele *spia-1(syb7920)*, bearing a deletion of 710 bp in *spia-1* with a modification of bp 789 (C -> T) to create a premature stop codon, and resulting in a truncated SPIA*-*1 protein of 26 aa (Fig 1C). Results obtained with *spia-1(syb7920)* phenocopied those obtained with *spia-1(fr179)* (Fig 1B and S1B). We further confirmed that the *spia-1*(*fr179)* mutation abrogates the PIA phenotype produced by RNA inactivation of any of the six furrow collagen genes (Fig 1D and S1C) but does not suppress the associated short size (i.e. the dumpy phenotype; Fig S1D). Moreover, we confirmed that *spia-1* mutation in furrow-less mutants reduced the endogenous expression of several AMPs genes including *nlp-29* (Fig S1E). Interestingly, we did not observe reduced *gst-4* nor *gdph-1* expression (Fig S1E), which are representative genes of the detoxification and osmotic stress responses kept in check by furrow collagens (Dodd et al., 2018). These data indicate that SPIA-1 is specifically required in the activation of the immune response upon furrow collagen loss.

In wild-type worms, *nlp-29* is expressed cyclically throughout development, possibly as a prophylactic protective mechanism following moulting (Aggad et al., 2023; Martineau et al., 2021; Sundaram and Pujol, 2024). We conducted a precise temporal analysis from the start of the L4 to the adult stage, using vulval shape as a proxy for developmental timing (Mok et al., 2015), as previously described (Aggad et al., 2023; Cohen et al., 2020). In the *nlp-29*p::GFP reporter strain, we observed a peak of GFP production at the L4.2-L4.3 and young adult stages (Fig 1E and 1F), suggesting a peak of *nlp-29* transcription a few hours before due to the folding time of GFP, which matches L3 & L4 moulting events respectively. A mutation in the transcription factor *sta-2* completely suppressed *nlp-29* induction, confirming that *sta-2* is required for the moulting-induced immune response (Fig 1E). In contrast, *spia-1(syb7920)* worms still presented both peaks and did not show any decrease in GFP production compared to wild-type worms (Fig 1E). In *dpy-7;spia-1(syb7920)* double mutants, although the PIA was reduced to levels qualitatively comparable to those in wild-type, as shown above, the moulting-induced peaks remained unaffected. Together, these results show that SPIA-1 is required in the furrow-less but not the moulting-induced immune responses.

### *spia-1* encodes a secreted protein with a novel cysteine-cradle domain

*spia-1* is predicted to encode a secreted nematode-specific protein of 165 aa with no previously known function (Davis et al., 2022). Its C-terminal region is annotated in the Panther database with a signature PTHR37435 that has no associated function (Davis et al., 2022; Thomas et al., 2022). This Panther family includes 5 other secreted *C. elegans* proteins: DPY-6, a mucin-like protein with a conserved role in cuticle formation (Sun et al., 2022), and 4 other uncharacterised nematode-specific proteins: F01G10.9, F13B9.2, Y34B4A.10 and F33D4.6 (Fig 2A). Orthologues of all 6 proteins are only present in nematodes, in free-living or in parasitic forms, and found in different clades, like *Rhabditina*, *Tylenchina* and *Spirurina*. These proteins have different lengths and contain regions with a compositional bias suggesting they may be largely intrinsically disordered, their common and most conserved part being located toward the C-terminus (Fig 2A and S2A). This common C-terminal region is predicted by AlphaFold2 (Jumper et al., 2021) to adopt a globular structure composed of four α-helices. These α-helices are arranged in two nearly orthogonal pairs, with two helices of one pair packed at both edges of the other and thus creating a cradle-shaped domain. This globular domain contains 4 invariant cysteines that define a sequence motif C1-(X)^22^-C2-(X)^7^-P-(X)^3^-C3- (X)^9^-C4. The cysteine residues are predicted to form two disulfide bonds connecting the α*-*helices, with C1 bonding C4 and C2 bonding C3 (Fig 2A, Sup movie). These disulfide bonds are likely to play a structural role and be essential for the maintenance of the cradle-like shape of the domain (Fig 2C and 2D). Owing to its features, this domain was named ’aECM cysteine-cradle domain’ (CCD-aECM or cysteine cradle domain) and its sequence diversity was added to the Pfam database (Mistry et al., 2021) as a new entry PF23626.

**Fig 2.**
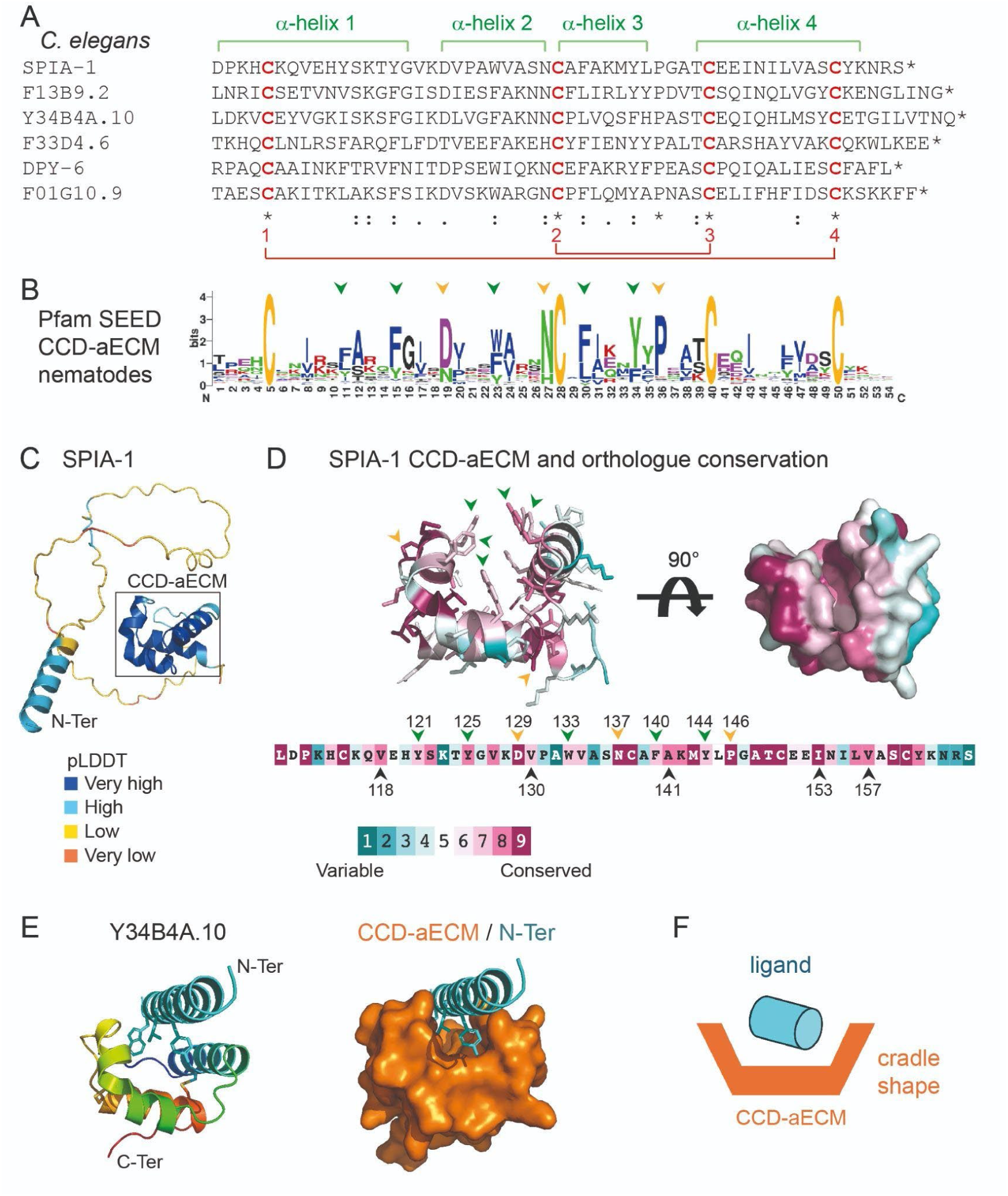
SPIA-1 is a secreted protein containing a novel cysteine-cradle domain. (A) In *C. elegans*, 6 proteins share a common and previously uncharacterised domain in their C-terminal region, of which the amino acid sequences are depicted. This domain contains 4 invariant cysteine residues predicted to form two disulfide bridges (red) connecting 4 α-helices (green) and was named the cysteine cradle domain (CCD- aECM). (B) The sequence logo derived from the Pfam (PF23626) SEED alignment shows residues of the domain conserved across nematode homologues. The relative size of the residue letters indicates their frequency in the aligned sequences of the Pfam SEED. Arrows point to the aromatic residues lining the groove (green) and other highly conserved amino acids (yellow). (C) SPIA-1 structural model predicted with AlphaFold2 (Abramson et al., 2024; Jumper et al., 2021), rendered in cartoon and coloured according to the Predicted Local Distance Difference Test score (pLDDT), which indicates how well a predicted protein structure matches protein data bank structure information and multiple sequence alignment data: dark blue >90, light blue <90 & >70, yellow <70 & >50, orange <50. The CCD-aECM domain is framed in black. (D) Amino acid sequence (bottom) and AlphaFold prediction of the CCD-aECM rendered in a cartoon, with the side-chains shown as sticks (top left) or in surface with a 90° rotation (top right), and coloured according to the Consurf conservation scores (Ashkenazy et al., 2016) based on SPIA-1 orthologs alignment. Arrows point to the aromatic residues lining the groove (green), aliphatic residues that are in contact (black), and other highly conserved amino acids (yellow). Numbers indicate the position of the amino acid in the full-length SPIA-1. The predicted structural model of SPIA-1 CCD-aECM is also shown on Sup Movie. (E) The AlphaFold prediction model of Y34B4A.10 (left) is rendered in cartoon and coloured in rainbow (blue to red indicating the path of the polypeptide chain from N- to C-terminal end). Residues from the α-helix that are predicted to engage in hydrophobic interactions are shown as sticks. The same model rendered in surface (right) demonstrates how the N-terminal α- helix of Y34B4A.10 (cyan) is predicted to bind to the CCD-aECM groove of this protein (orange). (F) Simplified illustration of the proposed model for the interaction of the CCD-aECM with a ligand.

The AlphaFold prediction of the cysteine cradle domain is in good agreement with predictions obtained using secondary structure and disulfide bond prediction programs which are based on different approaches (Buchan and Jones, 2019; Craig and Dombkowski, 2013; Drozdetskiy et al., 2015). The multiple sequence alignment of the cysteine cradle domain family (Fig 2D) or of the SPIA-1 orthologues in nematodes (Fig 2D) showed conserved features that strongly support the predicted structural model. In addition to the 4 invariant cysteine residues, these include: a highly conserved proline (Pro146 in SPIA-1) preceding and orienting the α-helix 4, thus facilitating disulfide bond formation; two highly conserved aspartate/asparagine (Asp129 & Asn137 in SPIA-1) at both caps of α-helix 2 needed for the sharp turns of the polypeptide chain. In addition, hydrophobic interior interactions between conserved aliphatic residues, as well as hydrogen bonds (e.g between Asn137 and the main- chain N-atom of Trp133 in SPIA-1), probably stabilise the cysteine cradle domain. The residues constituting the groove are also semi-conserved suggesting they may be important for function (Fig 2B and 2D, Sup Movie). Moreover, aromatic residues line the groove: a prominent tryptophan located on α-helix 2 (Trp133 in SPIA-1), a phenylalanine (Phe140 in SPIA-1) and 3 tyrosines (Tyr121, Tyr125, Tyr144 in SPIA-1); they define a highly hydrophobic interface that is probably involved in binding of unknown interaction partner(s). Interestingly, in the AlphaFold2 model of Y34B4A.10 (uniprot ID:Q8WSP0), an N-terminal α-helix of the protein itself docks into this groove (Fig 2E). A similar mode of interaction involving an α-helix docked into a hydrophobic groove has been previously observed; for example, between the p53 transactivation domain α-helix and the MDM2 cleft (Kussie et al., 1996). Alternatively, aromatic hydrophobic residues are also known to engage in binding of proline-rich peptides (Cottee et al., 2013; Macias et al., 2002) suggesting another potential functional interaction in which the cysteine cradle domain might be involved (Fig 2F).

*spia-1* is expressed in the epidermis and its expression oscillates during larval stages

During larval development, there are 4 moults spaced by ∼7-8 hours during growth at 25°C. Genome-wide transcriptomic studies have revealed the rhythmic activity of thousands of genes that align with the moulting cycle. They follow a repeated pattern of oscillations in each cycle, peaking at a distinct point in each larval stage. A large proportion of the cycling genes are expressed in the epidermis and are suggested to be required for the formation of the new cuticle (Hendriks et al., 2014; Kim et al., 2013; Meeuse et al., 2020; Meeuse et al., 2023; Tsiairis and Großhans, 2021). These oscillating transcripts include precuticle components that are only transiently present when the new cuticle is synthesised at intermolt and are endocytosed and degraded before each moult (Sundaram and Pujol, 2024). Analysing the data from (Meeuse et al., 2020; Meeuse et al., 2023), we observed that the transcripts for SPIA-1 and related proteins are part of these rhythmic oscillations (Fig 3A). We define that each cycle ended by the expression of AMPs, including *nlp-29*, which have been proposed to be induced to protect the epidermis while the old cuticle is shed (Martineau et al., 2021), and that one of the first genes to start to oscillate in the early L1 is *dpy-6*. It encodes a protein that, in addition to its CCD-aECM, is enriched with tandem repeats of serine and threonine residues similar to those found in highly glycosylated mucins (Sun et al., 2022). We then reanalysed the peak phase of genes that are known to be important for cuticle morphogenesis relative to *dpy-6*. All 5 genes encoding a CCD-aECM peak just after the pre-cuticle genes (orange) *let-653* and *fbn-1,* with *F01G10.9* (green) peaking together with the pre-cuticle genes *noah-1/2* & *lpr-3* and the 6 furrow Dpy collagens *dpy-2, dpy-3, dpy-7, dpy-8, dpy-9, dpy-10* (brown), followed by *spia-1, F13B9.2*, *F33D4.6* and *Y34B4A.10* (blue). These are then followed by the non-furrow collagens like *dpy-4, -5, -13* (black), and then the AMPs (pink) at the very end of each cycle (Fig 3B). The observation that CCD-aECM encoding genes cycle at the beginning of the new cuticle synthesis, together with precuticle and furrow collagen genes suggest a role in the formation of the new cuticle, including a very early role for *dpy-6*.

**Fig 3.**
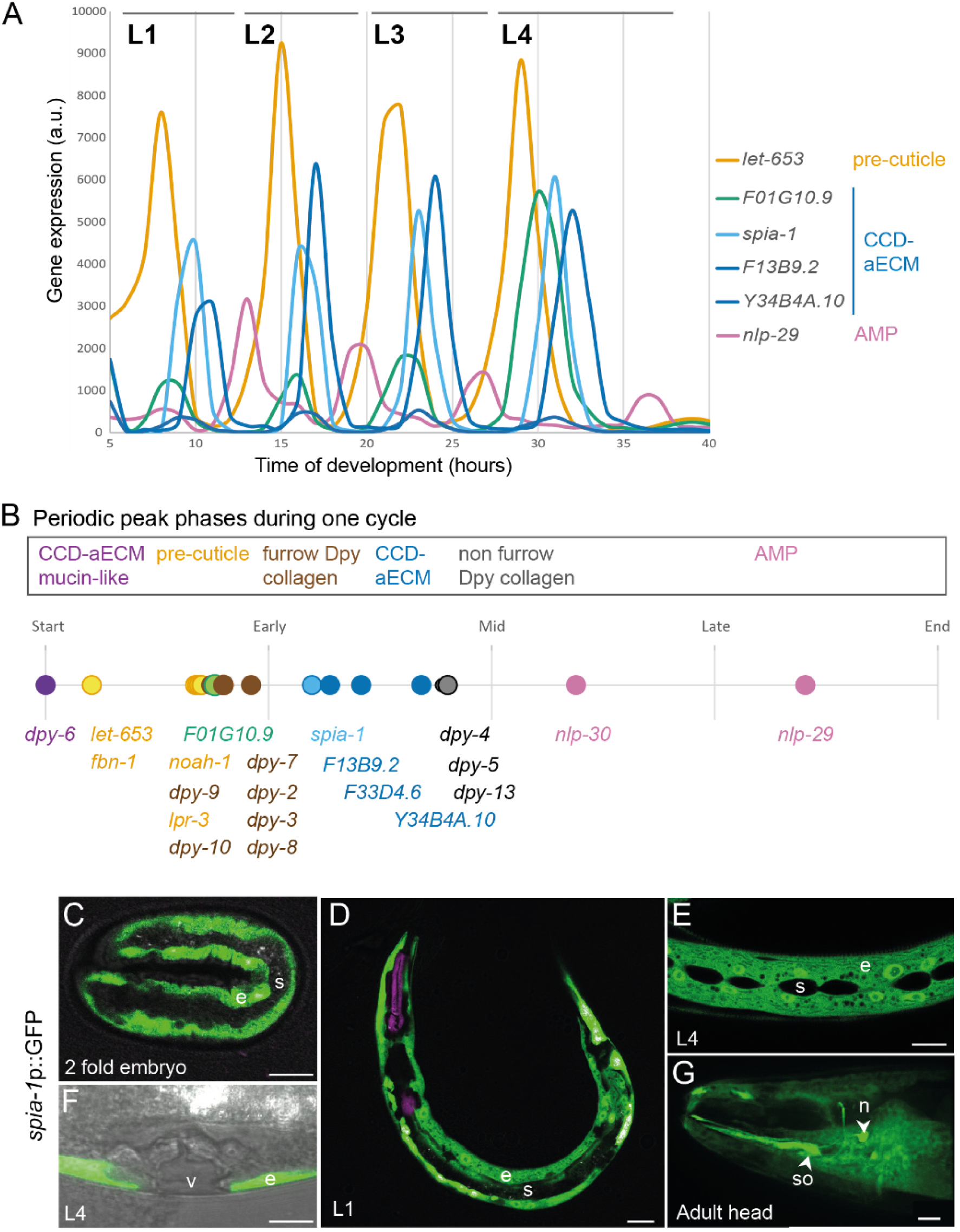
*spia-1* is expressed in the epidermis and oscillatory between each moult. (A) AMP and aECM gene expression oscillates between each moult, absolute levels of expression data from (Meeuse *et al*. 2020). (B) Between each moult, a timeline of gene expression is represented, with *dpy-6* starting each cycle. Data adapted from (Meeuse *et al*. 2020). (C-G) Expression pattern of *spia-1* transcript in worms carrying the *frEx631[pSO22(spia-1*p::GFP*), myo-2*p::mCherry] transgene. Representative confocal images, n>5, of (C) 2-fold embryo, (D) L1 larva, (E, F) L4 larva, and (G) adult head. The signal is visible in the epidermis (e) at all stages, and also in head socket cell (so) and a neuron (n), but not in the seam cell (s), nor the vulva (v). Grey colour in (F) was acquired with a transmitted detection module; scale bar, 10 µm.

Targeted DamID studies are consistent with SPIA-1 expression in the epidermis hyp7 in larval stages (Katsanos et al., 2021). In adult-specific single cell RNAseq, *spia-1* is also found expressed in cephalic and inner labial socket and phasmid sheath cells (Ghaddar et al., 2023). A transcriptional reporter confirmed that *spia-1* is expressed in the main epidermal cell and socket cells but is not visible in other epithelial cells like the seam cells, nor in the vulval cells (Fig 3C-G). The expression starts in embryos at the 2-fold stage, which is the time when pre-cuticle components like LPR-3 start to mark the cuticle, but later than the earliest components of the pre-cuticle sheath like NOAH-1 and FBN-1 (Fig 3C) (Balasubramaniam et al., 2023; Birnbaum et al., 2023; Cohen and Sundaram, 2020; Vuong-Brender et al., 2017). The transcriptional reporter might be missing some of the endogenous regulation, but it includes 1.2 kb of upstream genomic sequence that harbours several predicted binding motifs for transcription factors, including NHR-23 that is important for oscillatory gene expression in epithelial cells (Davis et al., 2022; Gerstein et al., 2010; Johnson et al., 2023). Taken together these observations suggest that *spia-1* is expressed in epidermal cells and oscillates with a peak phase that follows the six furrow collagens and precuticle components.

### SPIA-1 is localised to aECM periodic furrows

We tagged the SPIA-1 protein by insertion of GFP in 3 different positions: at the N-terminus after the signal peptide, at the C-terminus before the stop codon, or internally before the CCD-aECM. We introduced extra-chromosomal transgenes expressing these GFP-tagged SPIA-1 under the control of its own promoter into the *spia-1(syb7920)* mutant. Upon RNAi inactivation of *dpy-7*, the *spia-1(syb7920)* mutation suppressed the PIA phenotype; among the three transgenes, only the one containing the internally tagged SPIA-1::sfGFP (hereafter SPIA- 1::sfGFP-int) rescued the PIA phenotype (Fig 4B and 4C). This is consistent with the lack of observable GFP signal in either of the 2 other strains. In contrast, the functional protein SPIA- 1::sfGFP-int was visible in association with cuticle furrows starting in late embryonic stages and continuing throughout all larval stages (Fig 4D). We further generated a knock-in strain, IG2212, with SPIA-1 tagged with mNeonGreen (mNG) at the same position as the internal sfGFP. We confirmed that SPIA-1::mNG was located at furrows as it colocalised with DPY- 2::BFP (Fig S3B). In both strains, SPIA-1 was still present in the adult, but more faintly (Fig 4D and S3A). We could not detect it in the shed cuticle, but this could be due to its low signal. In addition, SPIA-1 accumulated in the vulva lumen during the mid-L4 stage (Fig 4D). In accordance with its cyclic expression, the fluorescence intensity of furrow-associated SPIA-1 also cycled, peaking in the middle of the L4 larval stage (Fig 4E). SPIA-1 strongly accumulated in vesicles preceding each moult, like precuticular components. During the L4 larval stage, the precuticular lipocalin LPR*-*3 was shown to be only transiently present, being secreted from the L4.3 stage onwards, and observed in an annular pattern between L4.4 to L4.7 (Forman-Rubinsky et al., 2017; Katz et al., 2022). Using vulval shape as a proxy for developmental timing, as described previously, we observed that in a double labelled mCherry::LPR-3, SPIA*-*1::sfGFP-int strain, SPIA-1 started to be visible at the L4.4 stage, at furrows in a pattern complementary to LPR-3, and in fluorescent vesicles (Fig 4E). Its level at the furrows then decreased, while it remained in vesicles throughout the end of the L4 stage.

**Fig 4.**
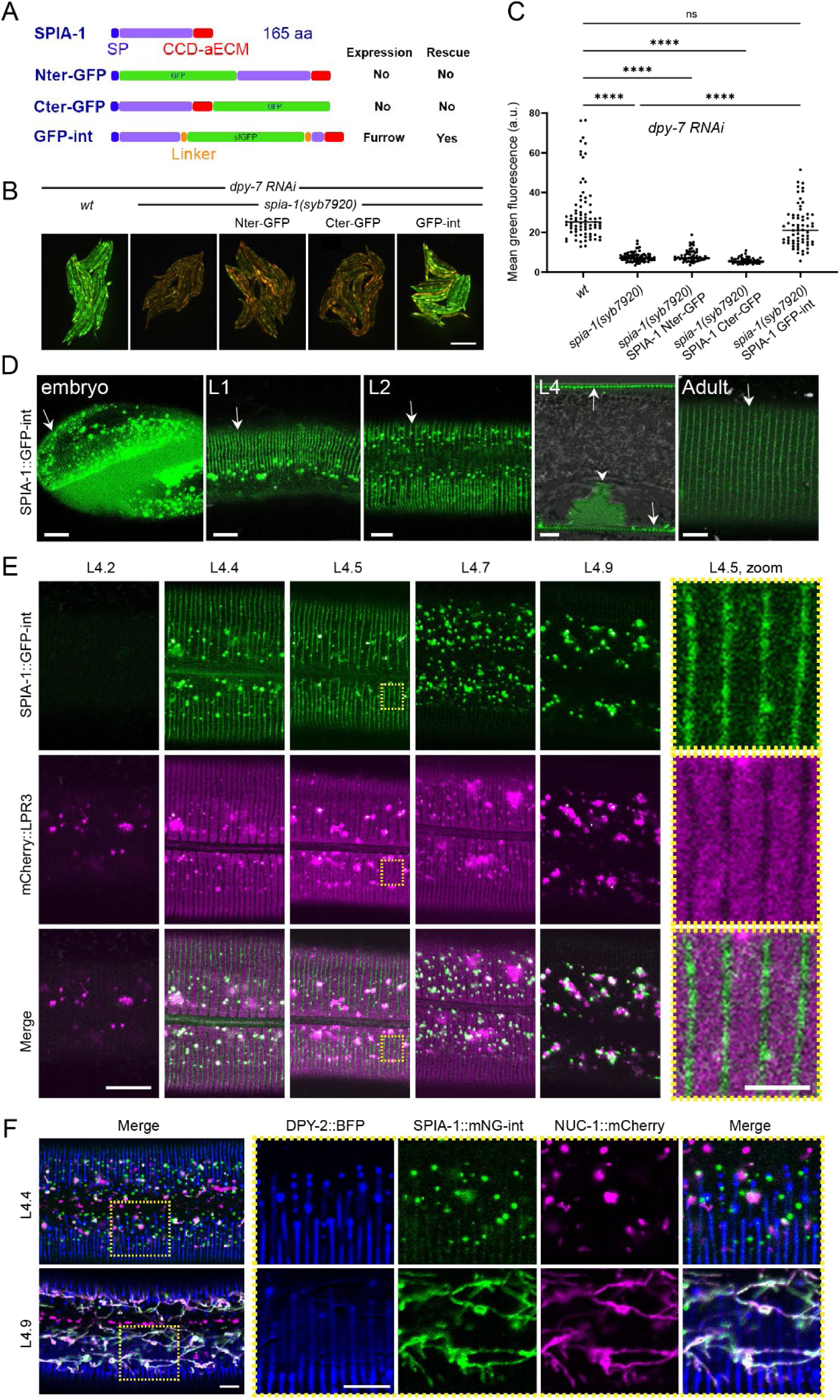
SPIA-1 localises to furrows. (A) Position of the insertion of GFP in each translational reporter with their expression pattern and rescue activities. For the KI strain, mNG is inserted at the same place as in GFP-int. (B-C) *spia-1* mutation suppresses *nlp-29*p::GFP overexpression in *dpy-7* worms. The rescue of this suppression has been tested in *spia-1(syb7920)* young adults with the extra-chromosomal gene producing SPIA-1 tagged with GFP in Nter, Cter or internal position, in three independent experiments. (B) Representative images of one experiment; scale bar, 500 µm. (C) Relative green fluorescence is quantified (n=58-79); \*\*\*\**p* < 0.0001. (D) Representative confocal images of the SPIA-1::sfGFP reporter (GFP-int) in 3-fold embryo, L1, L2, L4 vulval lumen and adult. We used a laser power ∼2 times higher in adults compared to other stages (see Fig S3A). White arrows and arrowhead indicate signal in furrows and in vulval lumen, respectively; n>5, scale bar, 5 µm. (E) The L4 stage is subdivided into sub-stages in relation to the shape of the vulva, as previously described (Mok et al., 2015). SPIA-1::sfGFP and mCherry::LPR-3 are observed in parallel. ∼7 times magnification of the areas contained in the dashed rectangles are provided on the far right; scale bar, 10 µm (left), 2 µm (magnified area). (F) Representative confocal images of L4.4 (top) and L4.9 (bottom) larvae expressing DPY-2::BFP, SPIA-1::mNG-int, and NUC-1::mCherry. ∼2.5 times magnification of the areas contained in the dashed rectangles are provided on the far right. Both single channels and the merge are shown; n>5, scale bar, 5 µm.

The intermolt peak and strong accumulation of SPIA-1 in vesicles before moulting resembled that of precuticular components like LPR-3 (Birnbaum et al., 2023; Forman- Rubinsky et al., 2017). Furthermore, we saw that SPIA-1::sfGFP-int and mCherry::LPR-3 vesicles partly overlapped (Fig 4E). To study further the nature of the SPIA-1 transient fluorescent vesicles, we combined SPIA-1::mNG with the lysosomal hydrolase NUC*-*1::mCherry that served as an endosomal and a lysosomal reporter, as previously described (Miao et al., 2020) and the DPY-2::BFP reporter to mark the furrows. Interestingly, in early L4, while NUC-1, SPIA-1 and DPY-2 are each visible in fluorescent vesicles, most of these were independent (Fig 4F). This suggests that SPIA-1 and DPY-2 are initially not associated with the same trafficking compartments, even though they colocalise at furrows once secreted in the matrix, and might be secreted in different ways. Remarkably, in late L4, most SPIA-1::mNG and NUC-1 fluorescent vesicles adopted a tubular structure, characteristic of lysosomal compartments, and appeared mostly colocalised (Fig 4F). It is interesting to note that these lysosomal tubular structures are visible with the mNG but not sfGFP tagged SPIA-1 strain, which presumably reflect the quenching of the latter’s fluorescence in acidic compartments. Overall, these data suggest that, in late L4, SPIA-1 is directed to lysosomes for degradation, a mark of precuticular components. This is also consistent with its reduced presence at furrows at the adult stage.

Altogether, these data suggest that SPIA-1 is an atypical cuticle component that shares some temporal and trafficking features of pre-cuticle, where its matrix signal peaks in the intermolt period of cuticle synthesis, after which most of it is cleared by endocytosis. Interestingly, the secreted hedgehog-related protein GRL-7, one of the few known components to be specifically positioned at the furrows in the pre-cuticle, contains a ground- like nematode-specific domain, which is another type of cysteine domain. It has been suggested to have a signalling role related to matrix association (Chiyoda et al., 2021; Serra et al., 2024; Sundaram and Pujol, 2024).

### SPIA-1 acts downstream of furrow collagens

*spia-1* was identified as a suppressor of the persistent immune activation provoked by the absence of furrows. Although *spia-1* did not reverse the Dpy phenotype of furrow-less mutants, it conceivably exerted its suppressive function by restoring normal furrow morphology. We examined the cuticle of worms deficient for different furrow collagens in the *spia-1(fr179)* background, with a COL-19::GFP marker and a DPY-7::sfGFP marker. In no case did the *spia-1* mutation restore furrows (Fig 5A and 5B). We noticed that when furrows are absent in a *dpy-2* mutant, the precuticle component LPR-3 cannot assemble anymore in its specific anti-furrow pattern during the mid L4, as visualised with sfGFP::LPR-3. The absence of *spia-1* could not restore the correct LPR-3 localisation in furrow-less mutants (Fig 5C). The absence of *spia-1* did not affect cuticle collagen DPY-7 or COL-19 nor precuticle LPR-3 localisation in an otherwise wild-type background (Fig 5A-C). Together, these observations indicate that *spia-1* acts downstream of the patterning and signalling roles of the furrow collagens.

**Fig 5.**
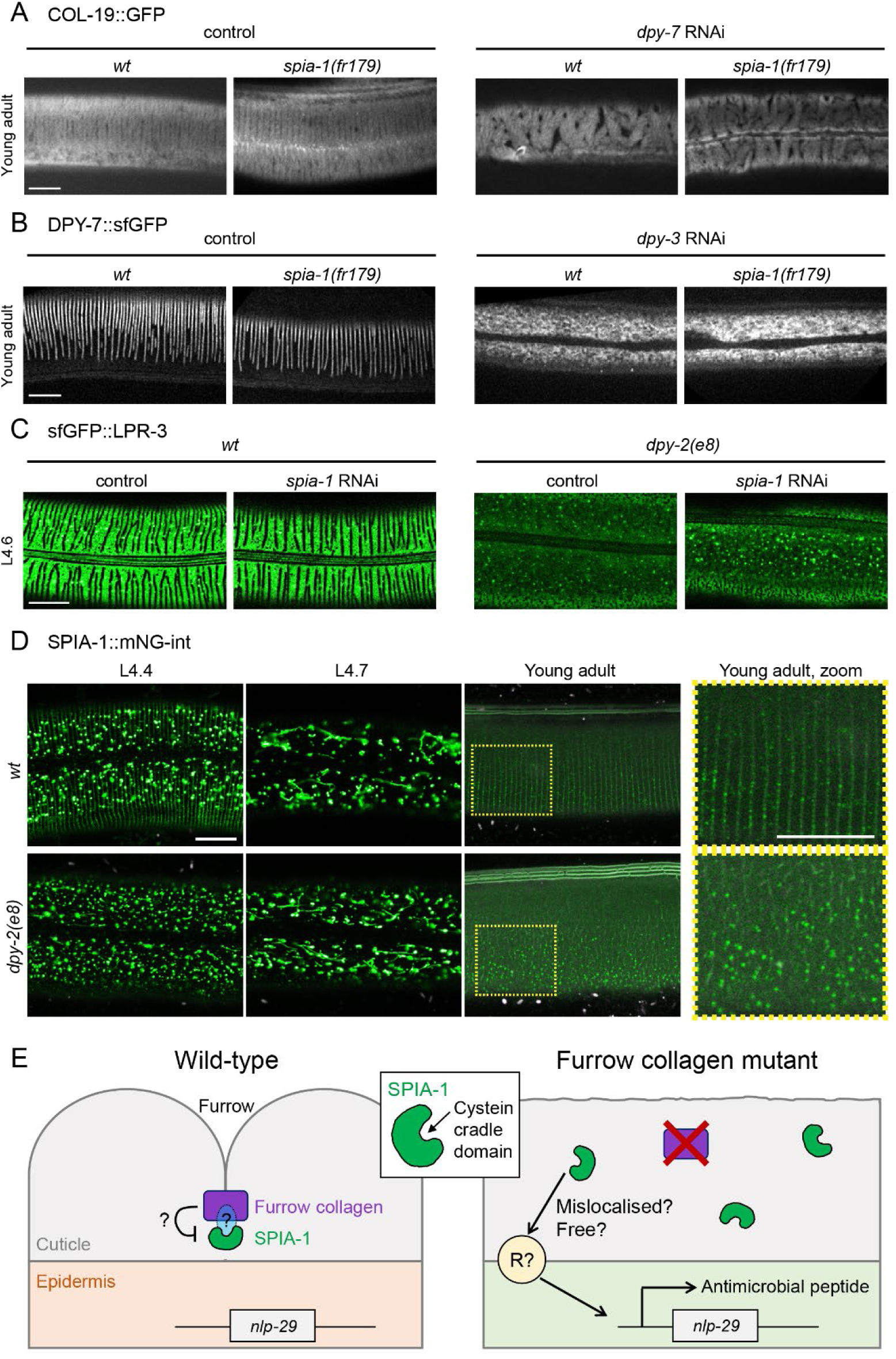
SPIA-1 acts downstream of furrow collagens. (A) *spia-1(fr179)* does not suppress COL-19::GFP abnormal pattern following *dpy-7* RNAi. Representative images of *wt* or *spia-1(fr179)* young adults carrying COL-19::GFP treated with *sta-1* or *dpy-7* RNAi bacteria; n>5, scale bar, 10 µm. (B) *spia-1* does not suppress the absence of furrows following *dpy-3* RNAi; representative images of *wt* or *spia-1(fr179)* worms carrying DPY-7::sfGFP treated with *sta-1* or *dpy-3* RNAi bacteria; scale bar, 10 µm. (C) *spia-1* does not suppress the abnormal sfGFP::LPR-3 in *dpy-2(e8)*; representative images of *wt* or *dpy-2(e8)* L4.6 worms carrying sfGFP::LPR-3 treated with control (*sta-1)* or *spia-1* RNAi bacteria; scale bar, 10 µm. (D) SPIA*-*1::mNG-int is mislocalised in *dpy-2(e8)* mutants. Representative images of *wt* or *dpy-2(e8)* L4.4, L4.7, or young adult worms carrying SPIA-1::mNG-int. We used a laser power ∼2 times higher in adults compared to other stages, see Fig S3A. A ∼2.5 times magnification of the areas contained in the dashed rectangles is provided on the far right; n>5, scale bar, 10 µm. (E) Cartoon presenting the proposed model for SPIA-1 activity in wild-type or furrow-less adults. Not to scale.

We then investigated SPIA-1 localisation in a furrow-less context, either in a *dpy-2* mutant or by RNAi inactivation of *dpy-7* (Fig 5D and S4). Furrow-less mutant conserved the vesicular and tubular pattern of SPIA-1 in L4 in the epidermis, and SPIA-1 was still present in the cuticle, suggesting that SPIA-1 was correctly produced, transported and degraded. However, in both L4 and adult, the furrowed pattern of SPIA-1 was lost and replaced by a signal randomly distributed in the cuticle (Fig 5D). We confirmed that the inactivation of any of the 6 furrow collagens, here *dpy-3*, impacts the localisation of other furrow collagens in the cuticle (Fig 5B) (McMahon et al., 2003). Thus, SPIA-1, like LPR-3, requires the presence of furrow collagens in the cuticle for its proper matrix localisation.

### A model for the role of SPIA-1 and its novel CCD-aECM domain

We propose a dual role of SPIA-1 as an atypical precuticular component of the furrow, as well as a cuticular sensor of cuticle damage. SPIA-1 shares characteristics of precuticular components as it is present at intermolt and highly endocytosed and degraded in lysosome before moulting. But unlike other characterised precuticular components, SPIA-1 is still present in the adult cuticle. Precuticular and some cyclic cuticular components like furrow collagens, are suggested to have a role in patterning the new cuticle ((Aggad et al., 2023; Forman-Rubinsky et al., 2017; Sundaram and Pujol, 2024), Fig 5A-C). In the absence of SPIA*-*1, we observed no obvious phenotype associated with cuticle morphogenesis, nor in patterning the precuticle LPR-3 or furrow collagens. Although this could mean that it has no role in cuticle morphogenesis, SPIA-1 could act redundantly, potentially with other cysteine cradle domain proteins. All the cysteine cradle domain proteins including SPIA-1 are only found in nematodes and predicted to be secreted. They are encoded by genes that have a cyclic expression and are predicted or already demonstrated to be expressed in the epidermis (Davis et al., 2022; Ghaddar et al., 2023; Katsanos et al., 2021; Meeuse et al., 2023). Altogether, these strongly suggest that all 6 cysteine cradle domain proteins are matrix proteins of the specialised nematode cuticle, hence the name of CCD-aECM for this novel Pfam domain.

SPIA-1 is required for the immune response provoked by the loss of furrows. In furrow- less mutants, we showed that SPIA-1 is aberrantly localised in the cuticle. The mislocalisation of SPIA-1 could trigger the persistent immune response in the epidermis. In normal circumstances, SPIA-1 could be muted by being directly or indirectly bound to collagen, thereby preventing it from signalling damage. In the absence of furrow collagen, it would be free to interact with unknown components, in parallel to the DCAR-1/GPA-12 pathway, to activate an immune response (Fig 5E). One hypothesis is that SPIA-1 could be directly linked to furrow collagens via its CCD-aECM. While 4 of the CCD-aECM proteins have not been studied yet, it is interesting to report that when endogenously tagged with mNG, DPY-6 is observed at furrows in the cuticle (Fig S5). Further investigations would be required to understand the potential role for CCD-aECM proteins in building a functional aECM and monitoring cuticle integrity.

## Materials and Methods

### EMS suppressor screen and mutation identification

P0 from the strain IG1389 *frIs7[nlp-29*p::GFP*, col-12*p::DsRed*] IV; frIs30[(col-19p::GPA-12gf), pNP21(pBunc-53::GFP)]* were mutagenised with EMS as previously described (Labed et al., 2012; Pujol et al., 2008a). Individual synchronised F2 L1 worms were plated on *dpy-7* RNAi plates. Late larval F2 that showed a low expression of *nlp-29*p::GFP were cloned. The ones that then showed a higher GFP expression at adult stage were further analysed. As a positive control for the mutagenesis, several candidates that still abrogate the GFP signal at the adult stage were also cloned, and led to the isolation of several new Nipi alleles including *snf-12(fr189)*. *spia-1* was further backcrossed with IG274 *frIs7* and a dozen of F2 without the GPA-12gf were isolated, then pooled to send their genome to sequencing (BGI). Sequences were analysed with MiModD v0.1.9, https://mimodd.readthedocs.io/en/latest, based on CloudMap (Minevich et al., 2012), using the sub-commands *varcall*, then *varextract* and finally *annotate*. This later step required SnpEff v5 and the SnpEff database *WBcel235.86*. The VCF file produced was re-formatted using a Python script to allow the curation of all putative mutations with user-defined thresholds.

### Nematode strains

All *C. elegans* strains were maintained on nematode growth medium (NGM) and fed with *E. coli* OP50, as described (Stiernagle, 2006). Table S1A shows a list of the strains used in this study, including those previously published: BE93 *dpy-2(e8) II* (Cox et al., 1980), IG1060 *sta-2(ok1860) V; frIs7[nlp-29p::GFP, col-12p::DsRed] IV* (Dierking et al., 2011), IG1689 *dpy-7(e88) X; frIs7[nlp-29p::GFP, col-12p::DsRed] IV* (Dodd et al., 2018), UP3666 *lpr-3(cs250[ssSfGFP::LPR-3]) X* & UP3808 *lpr-3(cs266[mCherry::LPR-3]) X* (Forman-Rubinsky et al., 2017), IG1389 *frIs7[nlp-29p::GFP, col-12p::DsRed] IV; frIs30[(col-19p::GPA-12gf), pNP21(pBunc-53::GFP)] I* & IG1392 *sta-2(ok1860) V; frIs7[nlp-29p::GFP, col-12p::DsRed] IV; frIs30[(col-19p::GPA-12gf), pNP21(pBunc-53::GFP)] I* (Lee et al., 2018), XW18042 *qxSi722[dpy-7p::DPY-7::sfGFP] II* & XW5399 *qxIs257[ced-1p::NUC-1::mCHERRY, unc-76(+)] V* (Miao et al., 2020), IG274 *frIs7[col-12p::DsRed, nlp-29p::GFP] IV; fln-2(ot611) X* (Pujol et al., 2008a), TP12 *kaIs12[COL-19::GFP]* (Thein et al., 2003), IG1426 *dcar-1(tm2484) V; frIs7[nlp-29p::GFP, col-12p::DsRed] IV; frIs30[(col-19p::GPA-12gf), pNP21(pBunc-53::GFP)] I* (Zugasti et al., 2014) and HW1371 *xeSi137[F33D4.6p:: gfp::h2b::pest::unc-54 3’UTR; unc-119 +] I* (Meeuse et al., 2023). Strains with extrachromosomal hygromycin resistance genes were selected on NGM plates supplemented with 0.3 mg/ml hygromycin B (Sigma-Aldrich).

### Constructs and transgenic lines

All following constructs were made using SLiCE (Motohashi, 2015) and the plasmid editor Ape (Davis and Jorgensen, 2022) and all primer sequences used to generate specific PCR amplicons are in Table S1B. A transcriptional construct (pSO22), was generated by cloning a PCR amplicon (3321-3322) containing 1.23 kb upstream of the *spia-1* start codon into pPD95.75 (Fire et al., 1990). To create translational constructs, GFP was inserted in *spia-1* either at the C-terminal ends (pSO24: 3326-3327) or after the N-terminal signal peptide (pSO25: 3366-3367, 2093- 3365). Internal tag translational construct (pSO26) was generated by inserting the sfGFP (kindly provided by A. Golden and H. Smith) in SPIA-1 at position 92 flanked with N-tag and C-tag linker used in pMLS288 and pMLS287 respectively (Schwartz and Jorgensen, 2016) (3392- 3393, 3394-3395).

pSO22 was injected at the concentration of 50 ng/µl with the co-injection marker *ttx-3*p::RFP at 50 ng/µl into N2 to get IG1986 and the co-injection marker *myo-2*p::mCherry at 2 ng/µl into N2 to get IG1988. Translational constructs (pSO24, pSO25, or pSO26) were injected at the concentration of 2 ng/µl, with the co-injection marker *myo-2*p::mCherry at 2 ng/µl and the HygR selection plasmid pZX13 at 50 ng/µl with pKS at 50 ng/µl into N2 (pSO24 and pSO25), or IG2093 *spia-1(fr179)* (pSO26) to get IG1999, IG2062, IG2108 respectively.

The strain PHX7920 *spia-1(syb7920)* generated by CRISPR editing (SunyBiotech), has a deletion of 710 bp (bp 79-788) in *spia-1* and a modification of bp 78 (C -> **<u>T</u>**) to create a premature stop codon. The sequence from the **<u>ATG</u>** to the original **<u>stop</u>** codon is <u>ATG</u>AAGCTAGTTGTTGTTTTGGCTTGTCTTGTTGTAGTAGCTGAGGCTTATTCAAAATCTGGAAATCC ATACAAGACT**<u>T</u>**AACTTGTGAGGAGATTAACATTTTGGTGGCCTCTTGCTACAAGAACAGAAGC**<u>TAA</u>**, resulting in a truncated SPIA-1 protein of 26 aa. All the transgenic strains carrying *spia-1(fr179)* or *spia-1(syb7920)* were obtained by conventional crosses and genotypes were confirmed by sequencing (see Table S1A for a list of all strains).

The strain IG2212 *spia-1(fr201(SPIA-1internal mNG^SEC^::3xFLAG))* was generated by CRISPR editing using a self-excising cassette as previously described (Dickinson et al., 2015); *mNG^SEC^::3xFLAG* was inserted at the same position than the GFP in pSO26. A repair template was constructed using Gibson cloning to insert a 622 bp 5’ homology arm and a 575 bp 3’ homology arm into an *Avr*II+*Spe*I digested pDD268 backbone (Dickinson et al., 2015) to make pJW2521. A sgRNA vector (pJW2568) targeting the ATCGGAAACAGTTGGTGGAG <u>TGG</u> sequence (PAM underlined, not included in vector) was made through SapTrap (Schwartz and Jorgensen, 2016), by cloning of an annealed oligo pair into pJW1839. Wild-type N2 animals were injected with pJW2568, pJW2521, a pCFJ2474 Cas9 plasmid, a *mlc-1*p::mNG co-injection marker (pSEM229), and a *snt-1*p::HisCl (pSEM238) counter selection marker (Aljohani et al., 2020; El Mouridi et al., 2020; El Mouridi et al., 2021). Plates were flooded with hygromycin and histamine as previously described (Dickinson et al., 2015). A hygromycin-resistant, rolling strain [JDW774 *spia-1((spia-1(spia-1 internal mNG^SEC^::3xFLAG)) X*] was recovered and then the self-excising cassette was removed through heat-shock as described in (Dickinson et al., 2015) to create IG2212. The strain MCP597 *dpy-2(bab597[DPY-2::mTaqBFP2]) II* was obtained by Segicel, by adding BFP at the C-terminus of DPY-2. The strain PHX3742 *dpy-6(syb3742(DPY-6::mNG))* was obtained by SunyBiotech, by adding mNG at the C-terminus of DPY-6. All knock-in strains were confirmed by PCR genotyping using primers outside the homology arms and Sanger sequencing.

### Sequence analyses

The following *C. elegans* CCD-aECM proteins (WormBase geneID/UniProt ID) SPIA-1/Q19281, Y34B4A.10/Q8WSP0, F33D4.6/O44189 (long isoform with the CCD-aECM), DPY-6/Q94185, F01G10.9/O17767 and F13B9.2/Q19385, were analysed with BlastP (Altschul et al., 1990), WormBase (Davis et al., 2022), Panther (Thomas et al., 2022), Pfam (Mistry et al., 2021), Interpro (Paysan-Lafosse et al., 2023) and AlphaFold2 & 3 (Abramson et al., 2024; Jumper et al., 2021). We built the Pfam family PF23626 (named ’aECM cysteine-cradle domain’) using sequences of CCD-aECM paralogues with domain boundaries defined based on the AlphaFold2 prediction models; Pfam PF23626 will be available in Pfam release 37.1. We iteratively searched for homologues using the HMMER package (Potter et al., 2018) and used an inclusion threshold of 27 bits. SPIA-1 orthologues including PIC17963.1 [*Caenorhabditis nigoni*], CAI5454296.1 [*Caenorhabditis angaria*], WKY17175.1 [*Nippostrongylus brasiliensis*], VDO70284.1 [*Heligmosomoides polygyrus*], EPB77628.1 [*Ancylostoma ceylanicum*], CDJ96309.1 [*Haemonchus contortus*], XP_013305412.2 [*Necator americanus*], KAF8381298.1 [*Pristionchus pacificus*], KAK5976273.1 [*Trichostrongylus colubriformis*], KAI6173309.1 [*Aphelenchoides besseyi*] were used for alignment and Consurf conservation scores (Ashkenazy et al., 2016).

### RNA interference

RNAi bacterial clones were obtained from the Ahringer library (Kamath et al., 2003) and verified by sequencing (see Table S1C). RNAi bacteria were seeded on NGM plates supplemented with 100 g/ml ampicillin and 1 mM Isopropyl-β-D-thiogalactopyranoside (IPTG). Worms were transferred onto RNAi plates as L1 larvae and cultured at 25 °C until L4 or young adult stage. In all our experiments, we use *sta-1* as a control, as we have shown over the last decade that it does not affect the development nor any stress or innate response in the epidermis (Dierking et al., 2011; Lee et al., 2018; Taffoni et al., 2020; Zhang et al., 2021; Zugasti et al., 2014).

### *nlp-29*p::GFP fluorescent reporter analyses

Representative fluorescent images including both green (*nlp-29*p::GFP) and red (*col- 12*p::DsRed) fluorescence were taken of *frIs7* transgenic worms mounted on a 2% agarose pad on a glass slide, anaesthetised with 1 mM levamisole in 50 mM NaCl, using the Zeiss AxioCam HR digital colour camera and AxioVision Rel. 4.6 software (Carl Zeiss AG). For quantification, the same worms were manually isolated and imaged again. Each worm was computationally contoured on ImageJ, by successively applying the RenyiEntropy threshold method provided by the plugin CLIJ2 (Haase et al., 2020) to the red image, converting the grayscale image to binary, suppressing noise (binary open), filling holes, and creating masks (analyze particles). Masks were applied to the original images, systematically controlled by eye, and corrected if needed. Mean green fluorescence signal was further measured for each contoured worm.

In figures S1C and S1D, *nlp-29*p::GFP expression was quantified with the COPAS Biosort (Union Biometrica; Holliston, MA) as described in (Labed et al., 2012). In each case, the results are representative of at least three independent experiments with more than 70 worms analysed. The ratio between GFP intensity and size (time of flight) is represented in arbitrary units.

### Confocal microscopy

Worms were mounted on a 2 % agarose pad, in a drop of 1 mM levamisole in 50 mM NaCl. Images were acquired during the following 60 min, using Zeiss confocal laser scanning microscopes (LSM780, 880 or 980) and the acquisition software Zen with a Plan-Apochromat Oil DIC M27 40×/1.4 or 63×/1.40 objective. Pinhole size was set to 1 AU. Samples were illuminated with 405 nm (BFP), 488 nm (GFP, mNG) and 561 nm (mCherry) with varied laser power based on protein abundance and tissue imaged, with 4 lines accumulation and 750 gain settings. Spectral imaging combined with linear unmixing was used to separate the autofluorescence of the cuticle.

### Quantitative PCR

Total RNA samples were obtained by Trizol (Invitrogen)/chloroform extraction. One mg of total RNA was then used for reverse transcription (Applied Biosystems). Quantitative real-time PCR was performed using 1 μl of cDNA in 10 μl of SYBR Green (Applied Biosystem) and 0.1 mM of primers on a 7500 Fast Real-Time PCR System using *act-1* as a reference gene. Primer sequences are provided in supplementary Table S1B.

### Statistical analysis

Data were analysed with the GraphPad Prism 10.3 software. Statistical differences between groups were determined by the Kruskal-Wallis’ test followed by the Dunn’s test. Data were considered significantly different when *p*-value was less than 0.05.

## Acknowledgements

We thank Damien Courtine for the installation and use of MiModD, Ebrima Bojang and Sarah Sharkaoui for participation in the genetic screen that identified *spia-1*, Matt Ragle, Emma Cadena, Jack Clancy for cloning and injections, Mathieu Fallet for help in image quantifications. We thank Andy Golden and Harold Smith for sending us the sfGFP plasmid, Margaux Gibert and SEGiCel (SFR Santé Lyon Est CNRS UAR 3453, Lyon, France) for the DPY- 2::BFP strain. The Fire Lab *C. elegans* Vector Kit was a gift from Andrew Fire (Addgene kit # 1000000001). Worm sorting was performed by Jérôme Belougne using the facilities of the French National Functional Genomics platform, supported by the GIS IBiSA and Labex INFORM. Some *C. elegans* strains were provided by the CGC, funded by NIH Office of Research Infrastructure Programs (P40 OD010440). We thank Tim Schedl and the staff at WormBase for amazing community work including the maintenance of a curated database, Meera Sundaram, Jonathan Ewbank, Michel Labouesse and Helge Großhans for discussions and comments on the manuscript. We thank the imaging core facility (ImagImm) of the Centre d’Immunologie de Marseille-Luminy (CIML) supported by the French National Research Agency through the “Investments for the Futureℍ program (France-BioImaging, ANR-10-INBS-0004).

## Supplementary Materials

**Fig S1.**
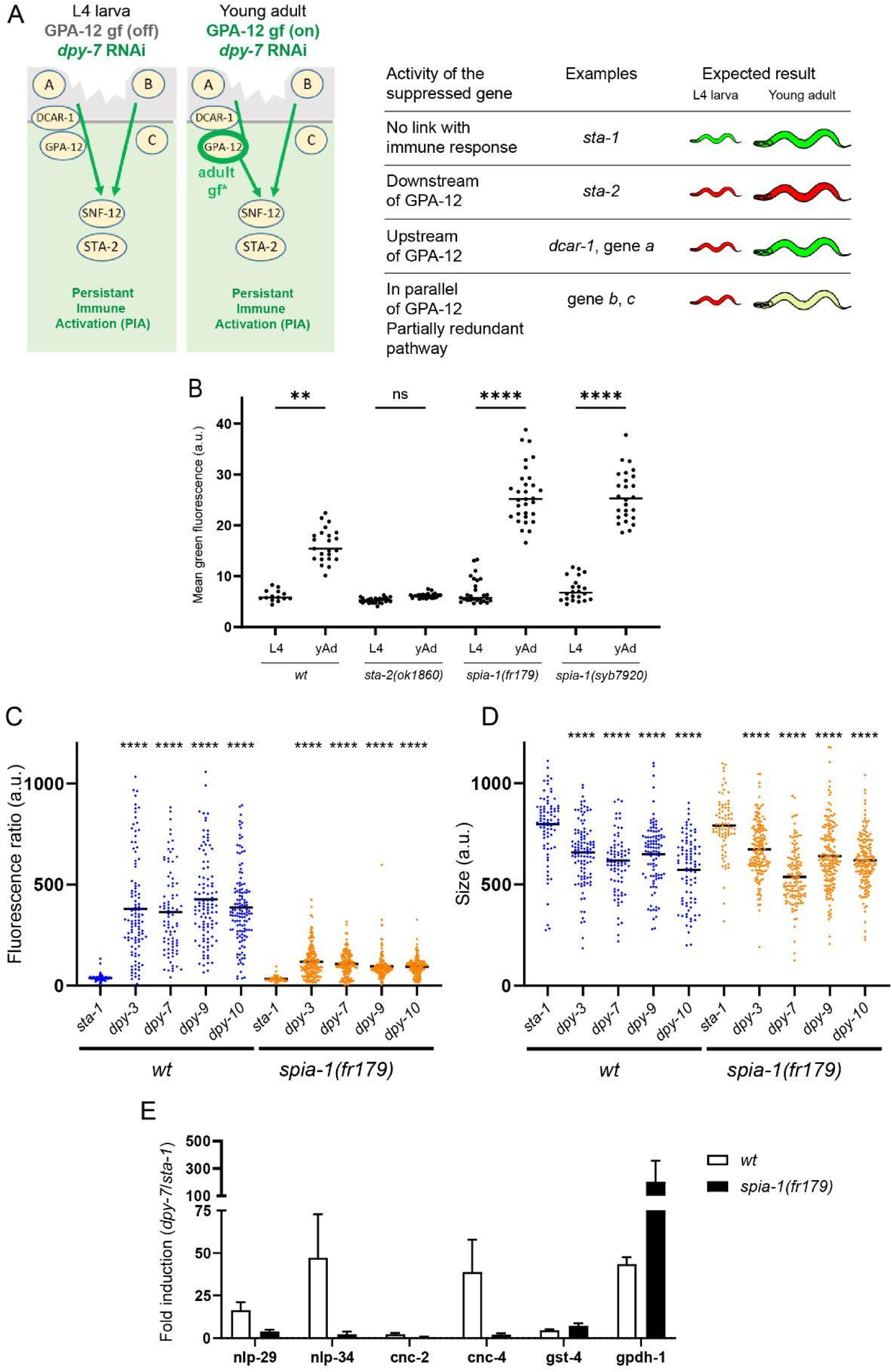
(A) In the suppressor screen, we triggered a PIA in the strain IG1389 by inactivating *dpy-7* by RNAi (left). In this strain, the state of the immune response is monitored (*frIs7* construct; green fluorescence off=inactive, green fluorescence on=active) and GPA-12 is constitutively active in the adult (*frIs30*). Different scenarios are expected depending on the gene affected after EMS-induced mutagenesis (right). (B) Quantification of relative green fluorescence in worms carrying *frIs7* and *frIs30* constructs, but without *dpy-7* RNAi inactivation, in L4 and young adults (yAd); n>14. Only the inactivation of a gene acting downstream of GPA-12 (e.g. *sta-2*) leads to the suppression of the green fluorescence in adults. (C-D) Quantification with the Biosort of the ratio between *nlp-29*p::GFP intensity and size (C) and of the size of the worms (D) in *wt* or *spia-1(fr179)* adults following RNAi inactivation of the 4 furrow collagen genes and the *sta-1* control; n>70, one of 3 independent experiments. *spia-1(fr179)* does not suppress the short size induced via inactivation of the 4 furrow collagen genes. Statistical comparisons were made by comparing to the corresponding *sta-1* control. \*\**p* < 0.01; \*\*\*\**p* < 0.0001. (E) mRNA levels of *nlp-29*, *nlp-34*, *cnc-2*, *cnc-4*, *gst-4* and *gpdh-1* were quantified by qPCR in wild-type and *spia-1(fr179)* worms upon RNAi inactivation of *sta-1* or *dpy-7*, in three independent experiments. The mean fold-changes between the *dpy-7* and *sta-1* levels are represented. In *spia-1(fr179)*, the transcription of AMPs genes including *nlp-29, nlp-34* and *cnc-4* were reduced, contrary to the transcription of *gst-4* and *gpdh-1*, the latter being increased.

**Fig S2.**
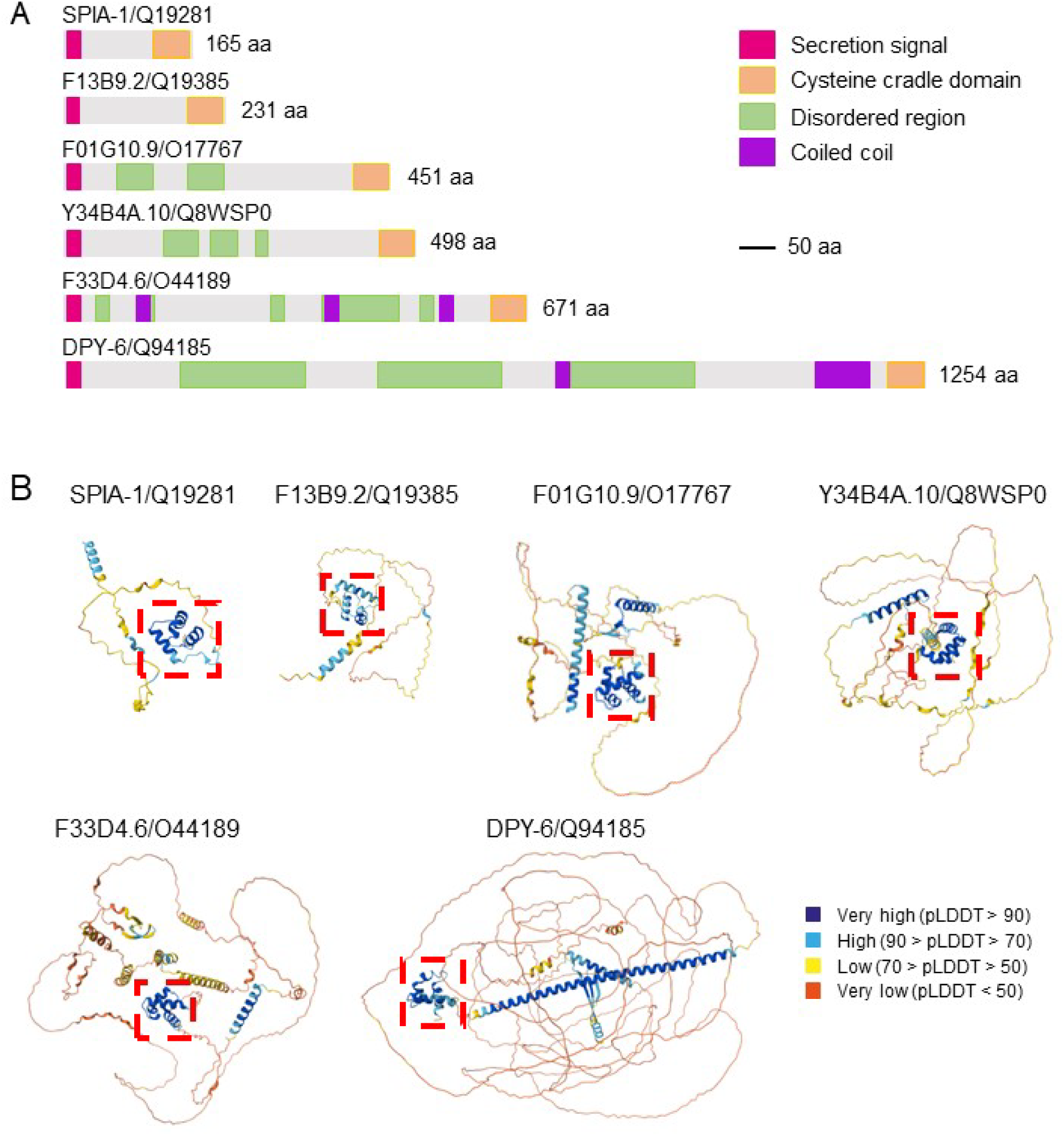
(A) Domain organisation of the 6 CCD-aECM proteins in *C. elegans*, as annotated in InterPro (Mistry et al., 2021; Paysan-Lafosse et al., 2023) and (B) structural models predicted with AlphaFold (Abramson et al., 2024; Jumper et al., 2021), rendered with the Predicted Local Distance Difference Test score (pLDDT), which indicates how well a predicted protein structure matches protein data bank structure information and multiple sequence alignment data.

**Fig S3.**
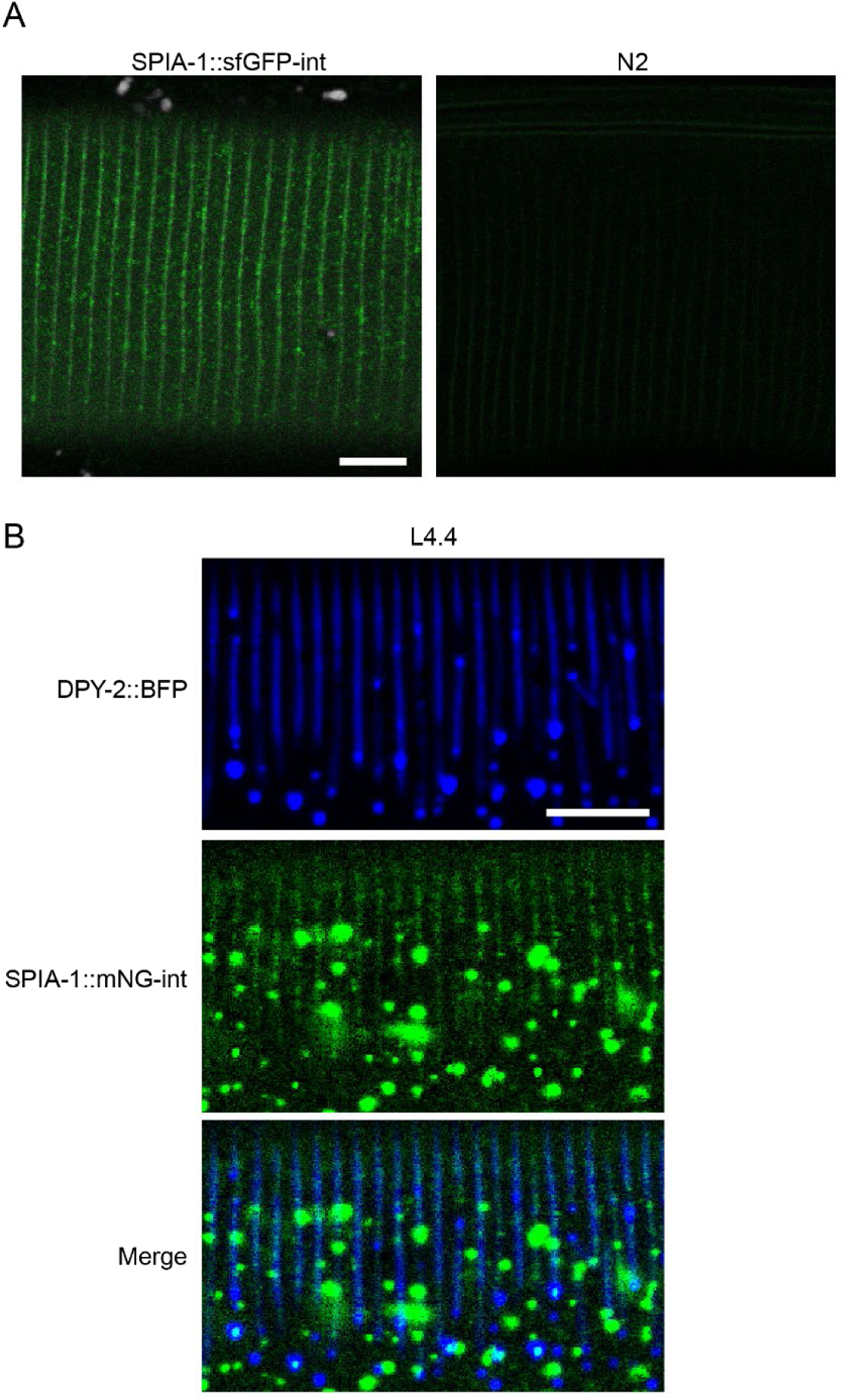
(A) The confocal image of the SPIA-1::sfGFP reporter (GFP-int) in the adult shown in figure 4D is presented aside from an adult N2 imaged using same illumination conditions; n>5, scale bar, 5 µm. (B) Zoom on the furrows in the L4.4 shown in Fig 4F. Both single channels and the merge are shown, as depicted. NUC-1::mCherry is not shown for clarity; scale bar, 5 µm.

**Fig S4.**
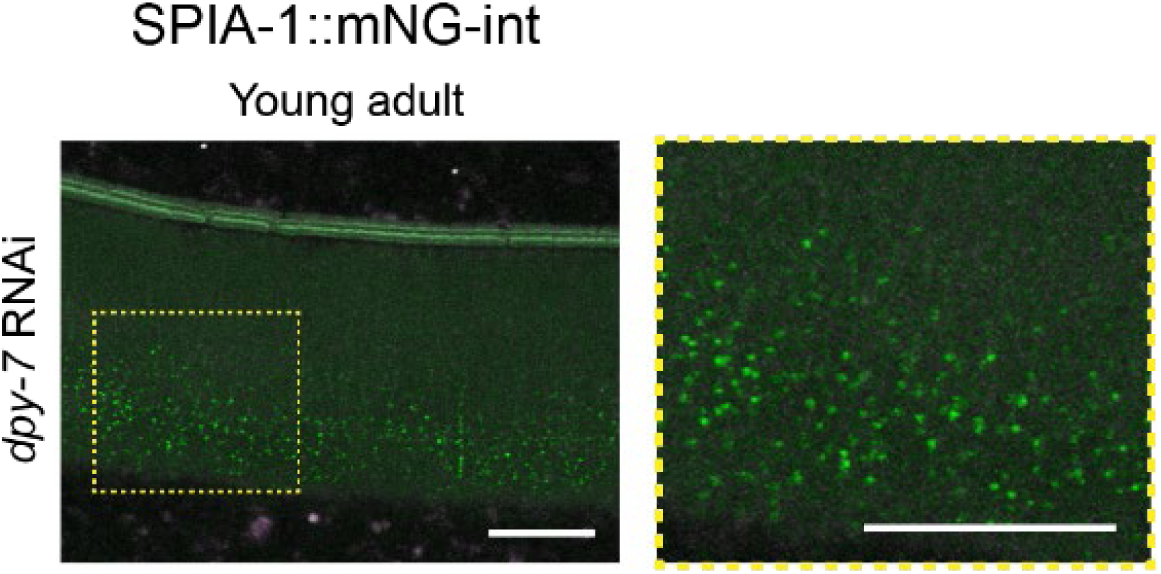
Representative images of SPIA-1::mNG-int young adults following *dpy-7* inactivation. To compare with Fig 5D. A ∼2.5 times magnification of the area contained in the dashed rectangle is provided on the far right; n>5, scale bar, 10 µm.

**Fig S5.**
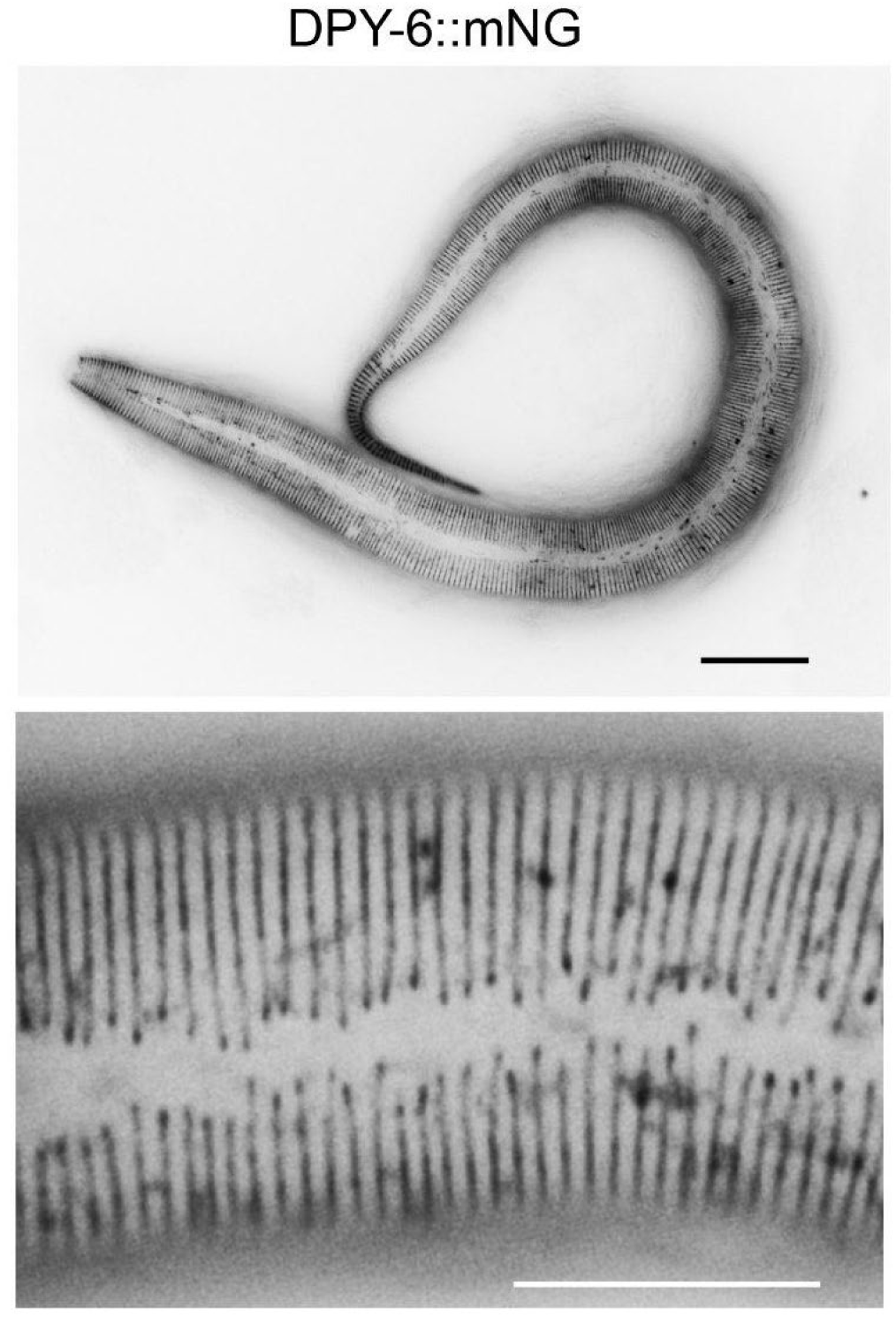
Representative fluorescent images of the furrow localisation of DPY-6::mNG-int, in a L1 (top) or L2 larva (bottom); n>5, scale bar, 20 µm (top), 10 µm (bottom).

## Supplementary movie

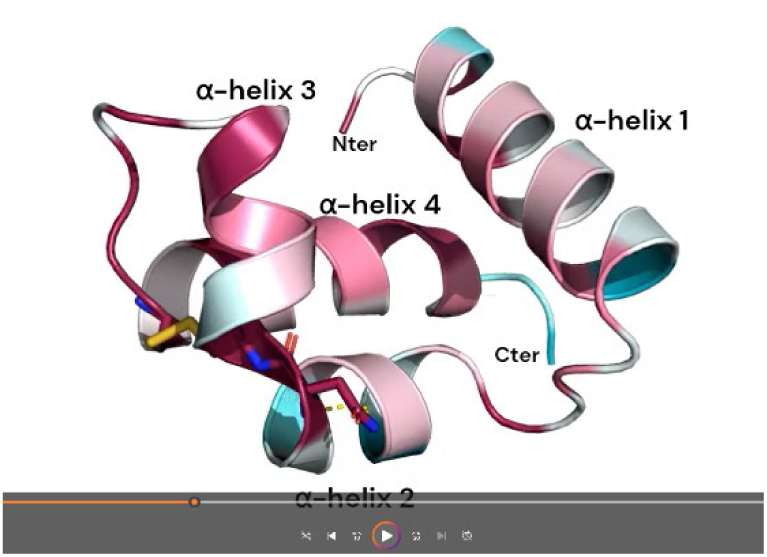

AlphaFold2 prediction of the SPIA-1 conserved CCD-aECM rendered in surface and cartoon. The model is coloured according to Consurf conservation scores across SPIA-1 orthologues in nematodes. The movie features the invariant cysteine residues predicted to form disulfide bonds (Cys138 with Cys150, Cys115 with Cys160), residues involved in hydrogen bonds probably stabilising the CCD-aECM (Trp133 with Asn137, Asp129 with Ala132, Leu110 with Ser159), aromatic residues lining the groove and defining a highly hydrophobic interface (Tyr121, Tyr125, Trp133, Phe140, Tyr144), and other conserved residues with predicted structural and functional roles (Gly126, Asp129, Pro146). Numbers indicate the position of the amino acid in the *C. elegans* SPIA-1 protein sequence. https://filesender.renater.fr/?s=download&token=5d52b645-c659-48da-abd3-291e7aae08dc

